# Future ocean warming may prove beneficial for the northern population of European seabass, but ocean acidification does not

**DOI:** 10.1101/568428

**Authors:** Sarah Howald, Louise Cominassi, Nicolas LeBayon, Guy Claireaux, Felix C. Mark

## Abstract

1.

The world’s oceans are acidifying and warming due to increasing amounts of atmospheric CO_2_. Thermal tolerance of fish much depends on the cardiovascular ability to supply the tissues with oxygen. The heart itself is highly dependent on oxygen and heart mitochondria thus might play a key role in shaping an organism’s tolerance to temperature. The present study aimed to investigate the effects of acute and chronic warming on respiratory capacities of European sea bass (*Dicentrarchus labrax* L.) heart mitochondria. We hypothesized that acute warming would impair mitochondrial respiratory capacities, but be compensated after long-term. Increasing *P*CO_2_ may cause intracellular changes, likely further constricting cellular energy metabolism.

We found increased leak respiration rates in acutely warmed heart mitochondria of cold-conditioned fish in comparison to measurements at their rearing temperature, suggesting a lower aerobic capacity to synthesize ATP. However, thermal acclimation led to increased mitochondrial functionality, e.g. higher RCR_o_ in heart mitochondria of warm-conditioned compared to cold-conditioned fish. Exposure to high *P*CO_2_ synergistically amplified the effects of acute and long-term warming, but did not result in changes by itself. We explained the high ability to maintain mitochondrial function under OA with the fact that seabass are moving between various environmental conditions. Improved mitochondrial capacities after warm conditioning could be due to the origin of this species in the warm waters of the Mediterranean. Our results also indicate that seabass are not yet fully adapted to the colder temperatures in their northern distribution range and might benefit from warmer temperatures.

**Summary statement:** Mitochondria of juvenile European sea bass hearts are impaired by acute warming, but seem to benefit from acclimation to warmer temperatures, they are only marginally impacted by ocean acidification.

**List of abbreviations**
Δ500: Acidification condition (ambient *P*CO_2_ + 500 µatm)
Δ1000: Acidification condition (ambient *P*CO_2_ + 1000 µatm)
A: Ambient *P*CO_2_ condition
C: Cold life conditioned group
CI: Complex I of the electron transport system
CII: Complex II of the electron transport system
CIV: Complex IV of the electron transport system
dph: Days post hatch
IPCC: Intergovernmental Panel on Climate Change
Leak-CI: State 4 respiration of complex I
MS-222: Tricaine methane sulfonate
OA: Ocean acidification
OAW: Ocean acidification and warming
OW: Ocean warming
OXPHOS: Full state 3 respiration of the electron transport system
OXPHOS-CI: State 3 respiration of complex I
OXPHOS-CII: State 3 respiration of complex II
OXPHOS-cytC: State 3 respiration after addition of cytochrome c
*P*CO_2_: Partial pressure of CO_2_
RCP: Representative concentration pathway
RCR_o_: Respiratory control ratio
State 4_o_: State 4 respiration after addition of oligomycin
W: Warm life conditioned group

## 3. Introduction

The increasing amount of atmospheric CO_2_ is working as a greenhouse gas raising atmospheric temperatures, and, as a consequence, also the sea surface temperatures (ocean warming – OW). At the same time, CO_2_ is partly taken up by the oceans, which leads to decreasing seawater pH (ocean acidification – OA). These processes together will lead to warmer and more acidic oceans, a trend that has already been observed over the last decades. Changes in water temperature have direct influence on the metabolic rate of ectothermic organisms, such as fish, with consequences for growth (Pörtner, et al., 2007; Peck, 2002), reproductive success (see review Llopiz, et al., 2014) and geographic abundance (Turner, et al., 2009; Pörtner, 2006). The body of studies looking into the effect of the changing environment on marine ectothermic organisms is growing rapidly. However, only a small part of these studies have investigated the effects of ocean acidification (OA) and the combined effects of ocean acidification and warming (OAW) on fish, with contrasting results between species as well as life stages (Kreiss, et al., 2015; Heuer & Grosell, 2014; Pope, et al., 2014; Bignami, et al., 2013; Frommel, et al., 2011). More investigation on a variety of fish species, on different life stages and under ecologically relevant *P*CO_2_ concentrations appears necessary (Pope, et al., 2014).

The effects of temperature on fish metabolism have been investigated intensively (e.g. Johnson & Katavic, 1986; Mirkovic & Rombough, 1998; Blásquez, et al., 1998; Farrell, 2002; Pörtner, et al., 2007; Hilton, et al., 2010; Strobel, et al., 2012). Thermal sensitivity of fish is mainly dependent on the capacity of the cardiovascular system to supply the tissues with oxygen (Pörtner & Lannig, 2009). The heart is a highly aerobic tissue (Driedzic, 1992) and it is therefore believed that the capacity of the heart mitochondria to produce ATP play a central role in defining thermal tolerance in fish. Although the subcellular processes are not yet fully understood, it has already been suggested that the functionality of heart mitochondria determines the temperature of heart failure (e.g. Chen & Knowlton, 2010; Iftikar & Hickey, 2013; Strobel, et al., 2013). Different mitochondrial processes have been shown to be impaired at elevated temperatures, which was indicated by decreased oxidative phosphorylation, decreased ATP production efficiency and lost integrity of the protein complexes of the electron transport system (e.g. Fangue, et al., 2009; Hilton, et al., 2010; Mark, et al., 2012; Iftikar & Hickey, 2013). Additionally, high temperatures increase the fluidity of mitochondrial membranes, potentially leading to increased proton leak through the inner mitochondrial membrane and decreased mitochondrial efficiency (Pörtner, 2012). Impaired mitochondrial metabolism thus might lead to alterations in cardiomyocyte ATP supply and consequently affect the performance of the cardiovascular system, which will ultimately determine the thermal sensitivity of the fish. Although juvenile and adult fishes generally possess well developed acid-base regulating mechanisms (see review Heuer & Grosell, 2014), increased ocean acidification may act as an additional stressor, e.g. by increased ATP demand for acid-base regulation. Consequently well-functioning mitochondria are even more important, if temperature and *P*CO_2_ are increased simultaneously. However, while the body of literature on the effects of temperature on mitochondrial function in fish is relatively large (e.g. Fangue, et al., 2009; Shama, et al., 2014 and literature therein), only a handful of studies investigated the effects of increased *P*CO_2_ on fish mitochondria and even less have combined OA and OW (but see Strobel, et al., 2013; Leo, et al., 2017). Yet as these two occur together, it will be important to determine their combined effects on mitochondrial metabolism to be able to take ecologically relevant conclusions. Furthermore, there is a relative lack of studies that focus on temperate, large and/or economically relevant species, while at the same time employing realistic *P*CO_2_ and exposure scenarios.

In our study, we exposed European seabass, *Dicentrarchus labrax* (L.), from 2 days post hatch (dph) for 7 months to three temperatures and two *P*CO_2_ conditions in a full-factorial design. Temperature and *P*CO_2_ conditions reflect the predictions of the IPCC for 2100 (IPCC, 2014). The European seabass is an important aquaculture species (160,000 t in 2015), but also an important target in commercial as well as in recreational fishing activities (Bjørndal & Guillen, 2018). European seabass are distributed throughout the Mediterranean, the Black Sea and the North-eastern Atlantic from Norway to Senegal in coastal waters from coastal lagoons and estuaries until 100 m depth (Bjørndal & Guillen, 2018). It has a relatively high tolerance to different temperatures and salinities (Via, et al., 1998; Claireaux & Lagardère, 1999). We studied the effect of OAW on mitochondria of juvenile seabass by determining mitochondrial capacities in permeabilized heart fibers of seabass juveniles that had been reared under the respective OAW conditions since hatch (7months). We examined effect of OAW on mitochondrial respiration rates of ATP producing processes (OXPHOS respiration) and the counteracting proton leak (Leak respiration), as well as their resulting ratio, the respiratory control ratio (RCR). An increased leak respiration, which is not compensated by increased OXPHOS respiration, results in a drop in RCR and indicates that mitochondrial functionality and consequently mitochondrial capacity to produce ATP is impaired.

We hypothesized that acute warming would impair mitochondrial performance in juvenile seabass hearts, as leak respiration may increase with thermal deterioration of mitochondrial membranes. On the other hand, compensational processes after long-term and developmental thermal acclimation could include changes in mitochondrial membrane properties, which would reduce leak respiration rates and consequently restore RCR. Additionally, we wanted to fathom the capacities of seabass mitochondria to cope with OA, especially when combined with OW. We hypothesized that the changes in intracellular *P*CO_2_ and bicarbonate concentration elicited by OA would affect mitochondrial metabolism, putting further pressure on the cellular energy metabolism.

## 4. Materials and Methods

The present work was performed within Ifremer-Centre de Bretagne facilities (agreement number: B29-212-05). Experiments were conducted according to the ethics and guideline of the French law and legislated by the local ethics committee (Comité d’Ethique Finistérien en Experimentation Animal, CEFEA, registering code C2EA-74) (Authorization APAFIS 4341.03, permit number 2016120211505680.v3).

All chemicals were purchased from Sigma-Aldrich, Germany, except for Tricaine methane sulfonate (MS-222), which was purchased from Pharma Q Limited, Hampshire, United Kingdom.

### 4.1. Animals and experimental conditions

#### 4.1.1. Animals

Larvae were obtained from the aquaculture facility Aquastream (Ploemeur-Lorient, France) at 2 dph (20.01.2016). Brood stock fish were caught in the sea off Morbihan, France. Four females (mean weight 4.5 kg) were crossed with ten males (mean weight 2.4 kg) which spawned naturally using photothermal manipulation. Conditions in the aquaculture facility during breeding were as follows: 13°C, 35 psu, pH 7.6, light conditions: 8h45m of light followed by 15h15m darkness. Spawning of eggs took place 15.01.2016; larvae hatched 18.01.2016 and were transported to our structures at 20.01.2016.

##### 4.1.1.1. Larval rearing

Larval rearing was performed in a temperature-controlled room using black 35 L tanks initially stocked with *ca*. 5000 larvae tank^−1^. The division into the experimental tanks took place at 3 dph (21.01.2016). During the following three days, the temperature for the warm condition was increased stepwise, 1°C during the first day and 2°C during each of the following days. The *P*CO_2_ conditions were applied directly after division into the experimental tanks. Starting at 7 dph (mouth opening), larvae were fed with live artemia, hatched from High HUFA Premium artemia cysts (Catvis, AE ’s-Hertogenbosch, Netherlands). Until 33 dph the artemia were fed to the larvae 24h after rearing cysts in sea water, afterwards the artemia nauplii themselves were fed with cod liver oil and dry yeast after 24 h and fed to the larvae after 48 h. The artemia were transferred to the larval rearing tanks from two storage tanks (one for each temperature) with peristaltic pumps, their concentration in the tanks was maintained high during the day, to allow *ad libitum* feeding, excess artemia left the tank via the waste water outflow. The 15 h photoperiod in the larval rearing room lasted from 7 am to 10 pm, the light intensity increased progressively during the larval rearing period from total darkness to 96 lux (Table S1). To work in the larval rearing facilities, headlamps were used (set to lowest light intensity). Larval mortality was 10-80 %, without pattern for condition (Table S2). Water surface was kept free of oily films using a protein skimmer. Density of larvae was continuously reduced for different samplings, such as growth or metabolic profiling. Additionally at approx. 750 dd, 300 larvae per tank were removed to decrease density in the tank, which corresponds to 38 dph and 49 dph for 20°C and 15°C rearing, respectively. At approx. 980 dd, the early juveniles were transferred from larval to juvenile rearing, corresponding to 50 dph and 65 dph for 20°C and 15°C rearing, respectively.

##### 4.1.1.2 Juvenile rearing

As they reached juvenile stage, fish were moved from the larval rearing facilities to juvenile tanks at approx. 1000 dd (50 dph, 08.03.2016 and 65 dph, 23.03.2016 for warm and cold life conditions, respectively). Fish were counted per tank and all fish from one condition were pooled in one tank until swim bladder test and separation into duplicate tanks at 1541 dd (78 dph, 05.04.2016) and 1301 dd (86 dph, 13.04.2016) for warm and cold life conditions, respectively. The swim bladder test was done to keep only the fish with developed swim bladders. Briefly, the fish were anaesthetized and introduced into a test container with seawater with a salinity of 65 psu. Those fish floating at the surface were removed from the test container and placed into the rearing tanks for recovery. The juveniles were reared in round tanks with a volume of 0.67 m^3^ and a depth of 0.65 m. Mortality rates of 24.8 to 43.4 % (Table S3) occurred between moving to juvenile condition and swim bladder test. During the first five days after moving to juvenile rearing, the juveniles were fed artemia nauplii (48h old and enhanced with cod liver oil and dry yeast) and commercial fish food. Commercial fish food (Neo Start) was fed daily and was adjusted in size (1-3) and amount during the juvenile rearing time, as recommended by the supplier (Le Gouessant, Lamballe, France). More precisely, they were fed *ad libitum* until 19.08.2016, from which food ratios were calculated after each sampling for growth, approx. every 3-4 weeks, using the formulae of Le Gouessant. The daily ration of food was supplied to the tanks by automatic feeders during day time; the fish were not fed during night time. Photoperiod was adjusted to natural conditions once a week, with slowly increasing light intensities in the juvenile rearing facilities during the first hour each morning. The tanks were cleaned daily after pH-measurements. Water flow within the tanks was adjusted once a week, so that oxygen saturation levels were not below 90%, with having equal flow through rates in all tanks of one temperature.

#### 4.1.2. Experimental conditions

The larvae and post-larval juveniles were reared under 6 different OAW scenarios, following the predictions of the Intergovernmental Panel on Climate Change (IPCC, (IPCC, 2014)) for the next 130 years. The acidification conditions included three different CO_2_ partial pressures (*P*CO_2_): today’s ambient situation in coastal waters of Brittany and the Bay of Brest (ambient group – A, approx. 650 µatm; cf. Pope, et al., 2014; Duteil, et al., 2016), a scenario according to the IPCC representative concentration pathway RCP6.0, projecting a Δ*P*CO_2_ of 500 µatm to current values (Δ500, approx. 1150 µatm) and a scenario according to RCP8.5, projecting a Δ*P*CO_2_ of 1000 µatm (Δ1000, approx. 1700 µatm). All acidification conditions were crossed with two different temperatures to create a ‘cold’ (C) and a ‘warm’ (W) life condition scenario: 15 and 20°C for the larval rearing followed by ambient temperature in the Bay of Brest (up to 18°C) and ambient plus 5°C for juvenile rearing, when ambient temperatures were exceeding 15°C. As larvae and post-larval juveniles would display different growth rates at the two different thermal scenarios, we adopted the concept of degree days (dph · T(°C)) as basis for comparison between these life conditions.

The sea water used in the aquaria was pumped in from the Bay of Brest from a depth of 20 m approximately 500 m from the coastline, passed through a sand filter (~500 µm), heated (tungsten, Plate Heat Exchanger, Vicarb, Sweden), degassed using a column, filtered using a 2 µm membrane and finally UV sterilized (PZ50, 75W, Ocene, France) assuring high water quality. During larval and early juvenile rearing, the water supply for the acidified incubation tanks came from a central header tank, where the water *P*CO_2_ conditions were adjusted. The water pH was controlled by an IKS Aquastar system (iks Computer Systeme GmbH, Karlsbad Germany), which continuously measured pH in one of the replicate tanks and opened a magnetic valve to bubble CO_2_ into the header tank when pH in the rearing tank became too high. Water exchange was set to 20 l/h until 12 dph and 25 l/h until the end of larval rearing. During juvenile rearing with higher water exchange rates, additional PVC columns were installed to control the pH in the rearing tanks. The water arrived at the top of the column and was pumped from the bottom of the column to the rearing tanks. The CO_2_-bubbling was installed at the bottom of the column and was adjusted by a flow control unit, when pH deviated from the desired value. One column supplied both replicate tanks of each condition. Temperature and pH were checked each morning with a handheld WTW 3110 pH meter (Xylem Analytics Germany, Weilheim, Germany; with electrode: WTW Sentix 41, NIST scale) before feeding the fish. The pH meter as well as the IKS Aquastar system were calibrated daily with NIST certified WTW technical buffers pH 4.01 and pH 7.00 (Xylem Analytics Germany, Weilheim, Germany). Total alkalinity was measured once a week following the protocol of Anderson & Robinson (1946) and Strickland & Parsons (1972): 50 ml of filtered tank water (200 µm nylon mesh) were mixed with 15 ml HCl (0.01 M) and pH was measured immediately. Total alkalinity was then calculated with the following formula:

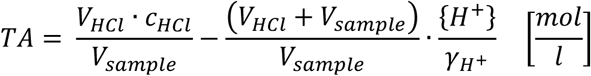

With: TA – total alkalinity [mol · l^−1^], V_HCl_ – volume HCl [l], c_HCl_ – concentration HCl [mol · l^−1^], V_sample_ – volume of sample [l], H^+^ – hydrogen activity (10^−pH^), γ^H+^ – hydrogen activity coefficient (here γ^H+^ = 0.758).

The Microsoft Excel macro CO2sys (Lewis & Wallace, 1998) was used to calculate seawater carbonate chemistry, the constants after Mehrbach et al. (1973, as cited in CO2sys) refit by Dickson and Millero (1987, as cited in CO2sys), were employed. Oxygen saturation (WTW Oxi 340, Xylem Analytics Germany, Weilheim, Germany) and salinity (WTW LF325, Xylem Analytics Germany, Weilheim, Germany) were measured once a week together with total alkalinity, from juvenile stage onwards, see all water parameters in Table 1.

**Table 1:** Water parameters during larval and juvenile phase of batch 2016: Larval period until 04.03.2016 (46 dph, ~900 dd) and 18.03.2016 (60 dph, ~900 dd) for warm and cold life conditions respectively, for the juveniles until 24.10.2016 (280 dph, ~5900 dd) and 08.02.2017 (387 dph, ~6200 dd) for warm-and cold-conditioned fish respectively. Means ± s.e.m. over all replicate tanks per condition. Temperature (Temp.), pH (free scale), salinity, oxygen (during juvenile rearing) and total alkalinity (TA) were measured weekly; *P*CO_2_ was calculated with CO2sys; sea water (SW) measurements were conducted in 2017 and 2018; A – Ambient *P*CO_2_, Δ500 – ambient + 500 µatm CO2, Δ1000 – ambient + 1000 µatm CO_2_, L – Larvae, J – Juveniles, C – cold life condition, W – warm life condition

| Treatment | pH<br>[-] | Temp.<br>[°C] | Salinity<br>[psu] | O2<br>[% airsat.] | TA<br>[mol l<sup>-1</sup>] | $PCO_2$<br>[$\mu\text{atm}$] |
| --- | --- | --- | --- | --- | --- | --- |
| L C A | 7.95±0.01 | 15.3±0.0 | 33.0±0.1 | - | 2364±17 | 656±16 |
| L C $\Delta 500$ | 7.77±0.01 | 15.3±0.0 | 33.0±0.1 | - | 2382±19 | 1041±26 |
| L C $\Delta 1000$ | 7.58±0.00 | 15.3±0.0 | 33.0±0.1 | - | 2394±26 | 1682±26 |
| L W A | 7.88±0.01 | 20.0±0.1 | 33.1±0.1 | - | 2369±21 | 832±13 |
| L W $\Delta 500$ | 7.79±0.01 | 20.0±0.1 | 33.1±0.1 | - | 2383±22 | 1057±30 |
| L W $\Delta 1000$ | 7.60±0.01 | 20.0±0.1 | 33.1±0.1 | - | 2380±23 | 1672±33 |
| J C A | 7.97±0.01 | 16.0±0.2 | 34.2±0.1 | 90.9±0.5 | 2396±18 | 655±18 |
| J C $\Delta 500$ | 7.75±0.01 | 16.0±0.2 | 34.2±0.1 | 92.2±0.6 | 2404±19 | 1107±21 |
| J C $\Delta 1000$ | 7.55±0.01 | 16.1±0.2 | 34.2±0.1 | 90.9±0.6 | 2399±19 | 1841±40 |
| J W A | 7.92±0.01 | 21.9±0.2 | 35.0±0.2 | 90.2±0.9 | 2418±12 | 788±22 |
| J W $\Delta 500$ | 7.78±0.01 | 21.8±0.2 | 35.0±0.2 | 90.5±0.7 | 2420±15 | 1133±43 |
| J W $\Delta 1000$ | 7.59±0.01 | 21.9±0.2 | 35.0±0.2 | 91.3±0.6 | 2423±12 | 1808±65 |
| SW cold | 8.05±0.01 | 14.5±0.5 | 33.0±0.2 | 101.2±0.6 | 2434±21 | 522±18 |
| SW warm | 7.95±0.02 | 21.2±0.4 | 32.7±0.1 | 102.3±1.4 | 2433±28 | 723±33 |

### 4.2. Mitochondrial respirometry

Measurements of mitochondrial respiratory capacities were done from approx. 3700 to 4100 dd in all conditions (183-199 dph and 234-249 dph in warm and cold life condition respectively). Although the same age in degree days was chosen, the cold-acclimated fish were significantly smaller than the warm-acclimated fish (length: 8.72±0.09 cm and 9.59±0.09 mm, lsmeans ± se, p<0.001, respectively and carcass weight: 10.00±0.45 g and 13.38±0.46 g, respectively, lsmeans ± se, p<0.001, LME), resulting in smaller ventricle sizes (0.0105±0.0004 g and 0.0122±0.005 g, respectively, p<0.05, lsmeans ± se, LME), as well as lower hepatosomatic indices(1.51±0.05 and 2.45±0.06, respectively, lsmeans ± se, p<0.0001, LME). However, condition factor was not different in the two temperature conditions (1.51±0.01 and 1.48±0.01, respectively, lsmeans ± se, p>0.1, LME), see further Table S4. *P*CO_2_ did not have an effect on fish size. Prior to the experiments, the fish were starved for two days. Two batches of eight fish each were processed per day. Juveniles were randomly caught from their tank and anesthetized with MS-222. The concentration of the anesthetic was adjusted to reach a loss of equilibrium within less than 5 minutes, typically 0.2 g l^−1^. Weight, fork length and body length were directly determined with a precision balance (Mettler, Columbus, Ohio, USA) and a caliper, to the next 0.01 g and 0.01 mm, respectively. Afterwards fish were killed by a cut through the neck, and the heart was completely dissected from the fish followed by excavation of the ventricle. Excess blood was removed from the ventricle by cleaning it on blotting paper, prior to weighing (Sartorius, Göttingen, Germany). The whole ventricle was used for respiration measurements. The ventricle was then stored in relaxing and biopsy preservation solution BIOPS (10 mM Ca-EGTA (0.1 µM free calcium), 20 mM imidazole, 20 mM taurine, 50 mM K-MES, 0.5 mM DTT, 6.56 mM MgCl_2_, 5.77 mM ATP, 15 mM phosphocreatine, pH 7.1 modified after Gnaiger, et al., 2000)) until all eight ventricles were dissected. The ventricles were then manually frayed and were permeabilized on ice with saponin (50 µg ml^−1^) for 20 min on a shaking table (80 rpm), followed by two cleaning steps for 10 min at 80 rpm in modified mitochondrial respiration medium MiR05 (160 mM sucrose, 60 mM K-lactobionate, 20 mM taurine, 20 mM HEPES, 10 mM KH_2_PO_4_, 3 mM MgCl_2_, 0.5 mM EGTA, 1 g l^−1^ bovine albumin serum (fatty acid free), pH 7.45 at 15°C, modified after Fasching, et al., 2014). During the permeabilization step, the livers and the carcasses of the fish were weighed to calculate hepatosomatic index (HSI) and condition factor (K), see Table S4.

Mitochondrial respiration of the permeabilized heart fibers was measured using four Oroboros Oxygraph-2K respirometers with DatLab 6 software (Oroboros Instruments, Innsbruck, Austria). Permeabilized fibers have the advantage of resembling the living state as closely as possible, while still allowing to control the supply of substrates and inhibitors to the mitochondria (Saks, et al., 1998; Pesta & Gnaiger, 2012). Measurements were conducted at 15°C and 20°C for all treatments to determine the effect of acute temperature changes on mitochondrial metabolism *in vitro*. The oxygen sensors were calibrated in air-saturated buffer prior to the experiments and in oxygen depleted buffer after each experiment. The measurements were done in MiR06 buffer (MiR05 buffer enriched with 280 u/ml catalase, (Fasching, et al., 2014)) to allow for reoxygenation with H_2_O_2_. A substrate-uncoupler-inhibitor titration protocol was employed to measure the capacities of the different complexes: Glutamate (10 mM), malate (2 mM) and pyruvate (10 mM) were used to measure state 4 respiration of complex I (Leak-CI, Table 2), followed by addition of ADP (2.5 mM) to measure state 3 respiration of complex I (OXPHOS-CI). Succinate (10 mM) and further ADP (twice 2.5 mM) resulted in full state 3 respiration (OXPHOS). Cytochrome c (0.01 mM) was used as control for inner mitochondrial membrane integrity (OXPHOS-cytC); measurements with increases of more than 10% following cytochrome c addition were not used for further analyses. Oligomycin (4 µg/ml) was used to inhibit F_O_F_1_-ATPase, resulting in state 4_oligomycin_ respiration (state 4_o_). Stepwise titration of FCCP (0.05 µl of 2mM stock solution per step) was used to uncouple the mitochondrial electron transport system and determine its maximum capacities. After uncoupling, complex I (CI), II (CII) and III were successively inhibited with rotenone (0.005 mM), malonate (5 mM) and antimycin A (0.0025 mM), respectively. Residual respiration after antimycin A addition was used to correct all mitochondrial respiration rates. Complex IV (CIV) capacities were then determined by addition of ascorbate (2mM) and TMPD (0.5 mM). Oxygen levels were usually restored by addition of 2 µl H_2_O_2_ (3% stock solution) after addition of oligomycin and again after addition of rotenone.

**Table 2:** Analyzed mitochondrial metabolic states (after Nicholls & Ferguson, 2002) during the substrate uncoupler inhibitor titration protocol of this study. Oxygen is available in all these states.

| Description | State | Name |
| --- | --- | --- |
| Substrates for CI are available, no ADP | 4 | Leak-CI |
| Substrates for CI and ADP are available and fuel state 3 respiration of CI | 3 | OXPHOS-CI |
| Substrates for CI and CII as well as ADP is available to fuel state 3 respiration of the electron transport system | 3 | OXPHOS |
| Oligomycin inhibits F<sub>0</sub>F<sub>1</sub>-ATPase resulting in state 4<sub>oligomycin</sub> respiration | 4<sub>o</sub> | state 4<sub>o</sub> |
| State 3 respiration of CII was calculated: OXPHOS - OXPHOS-CI | 3 | OXPHOS-CII |
| Respiratory control ratio was calculated: OXPHOS · state 4<sub>o</sub><sup>-1</sup> | - | RCR<sub>o</sub> |

A measure for state 3 respiration of CII (OXPHOS-CII) was calculated (OXPHOS-CII = OXPHOS – OXPHOS-CI), although this measure will lead to lower respiration rates than determination of state 3 respiration, when only substrates for CII are available (Gnaiger, 2009; Mark, et al., 2012). For all complexes, the fraction of the respiration of the respective complex on OXPHOS respiration was calculated. This was also done for state 4_o_, which we tentatively termed ‘state 4_o_-fraction’, as a relative indicator of proton leak, despite the fact that membrane potential is higher in state 4_o_ than in state 4. Mitochondrial quality and efficiency was evaluated by calculating the respiratory control ratio (RCR_o_) (OXPHOS · (state 4_o_)^−1^), which is an indicator of mitochondrial coupling (Gnaiger, 2009; Strobel, et al., 2013).

### 4.3. Statistical analysis

All statistics were performed with R (Core-Team, 2017): Data were tested for outliers (Nalimov test), normality (Shapiro-Wilk’s test, p > 0.05) and homogeneity (Levene’s test, p > 0.05). Mitochondrial respiratory data were fitted to linear mixed effect models (LME, “lme” function of “nlme” package, (Pinheiro, et al., 2017)). Acclimation and assay temperature, as well as *P*CO_2_ concentrations and their interactions were included as fixed effects. Due to the significantly different sizes of the fish, fish weight was also included as fixed effect, whereas the oxygraph chamber and the number of the run at that day were included as random effects. In case of heterogeneity of data, variance structures were included in the random part of the model; the best variance structure was chosen according to lowest Akaike information criteria (AIC) values. Validity of linearity for *P*CO_2_ concentration and weight was cross tested with generalized additive models (“gam” function of “mgcv” package, (Wood, 2017)), as described in Zuur et al. (2009). If linearity was given, the linear mixed effect model was chosen instead of the generalized additive model. If significant effects were detected in the linear mixed effect models, posthoc Tukey tests were performed with the “lsmeans” function (“lsmeans” package, Lenth, 2016). Significance for all statistical tests was set at p < 0.05. All graphs are produced from the lsmeans-data with the “ggplot2” package (Wickham, 2016). All data are shown as lsmeans ± s.e.m‥ Biometrical data were also tested with LME models, with *P*CO_2_, acclimation and assay temperature as fixed effects and origin tank as random effect. Model validation was done as described for mitochondrial respiratory data.

## 5. Results

Acclimation and assay temperature had effects on a number of the investigated capacities of mitochondrial complexes and properties. On the other hand *P*CO_2_ was only significant in interaction with acclimation or assay temperature, not as a factor itself. Fish mass only had significant effects on RCR_o_ and CIV respiration rates. Additionally, in some of the other investigated mitochondrial complexes and properties trends with fish mass were visible. F-values and significance of the fixed effects from the LME models on the different mitochondrial oxygen consumption rates and their ratios to OXPHOS respiration are presented in Table 3.

**Table 3:** F-values of fixed effects from the linear mixed models on mitochondrial respiration of permeabilized heart fibers of juvenile European seabass; Den. d.f. – Denominator degrees of freedom, no d.f. – number degrees of freedom, AT – acclimation temperature, MT – assay temperature.

|  | Den. | AT | MT | PCO<sub>2</sub> | Weight | AT:MT | AT:PCO<sub>2</sub> | MT:PCO<sub>2</sub> | AT:MT:PCO<sub>2</sub> |
| --- | --- | --- | --- | --- | --- | --- | --- | --- | --- |
|  | d.f. | no d.f. 1 | no d.f. 1 | no d.f. 2 | no d.f. 1 | no d.f. 1 | no d.f. 2 | no d.f. 2 | no d.f. 2 |
| RCR<sub>o</sub> | 126 | 12.50<sup>***</sup> | 1.66<sup>n.s.</sup> | 0.19<sup>n.s.</sup> | 4.73<sup>*</sup> | 1.31<sup>n.s.</sup> | 2.18<sup>n.s.</sup> | 0.05<sup>n.s.</sup> | 0.72<sup>n.s.</sup> |
| Oxygen consumption rates |  |  |  |  |  |  |  |  |  |
| OXPHOS | 147 | 40.83<sup>***</sup> | 20.76<sup>***</sup> | 2.46<sup>·</sup> | 0.65<sup>n.s.</sup> | 0.09<sup>n.s.</sup> | 2.16<sup>n.s.</sup> | 2.04<sup>n.s.</sup> | 4.20<sup>*</sup> |
| OXPHOS-CI | 146 | 29.08<sup>***</sup> | 10.10<sup>**</sup> | 1.17<sup>n.s.</sup> | 1.09<sup>n.s.</sup> | 0.00<sup>n.s.</sup> | 3.33<sup>*</sup> | 0.20<sup>n.s.</sup> | 3.46<sup>*</sup> |
| OXPHOS-CII | 144 | 12.30<sup>***</sup> | 26.30<sup>***</sup> | 2.05<sup>n.s.</sup> | 0.22<sup>n.s.</sup> | 0.00<sup>n.s.</sup> | 0.15<sup>n.s.</sup> | 4.14<sup>*</sup> | 3.52<sup>*</sup> |
| State 4<sub>o</sub> | 141 | 122.51<sup>***</sup> | 39.75<sup>***</sup> | 1.78<sup>n.s.</sup> | 2.87<sup>·</sup> | 5.43<sup>*</sup> | 4.03<sup>*</sup> | 0.93<sup>n.s.</sup> | 0.18<sup>n.s.</sup> |
| CIV | 129 | 38.64<sup>***</sup> | 42.90<sup>***</sup> | 0.47<sup>n.s.</sup> | 10.83<sup>**</sup> | 2.82<sup>·</sup> | 0.57<sup>n.s.</sup> | 0.08<sup>n.s.</sup> | 0.15<sup>n.s.</sup> |
| Fractions of OXPHOS respiration |  |  |  |  |  |  |  |  |  |
| CI | 144 | 2.93<sup>·</sup> | 0.56<sup>n.s.</sup> | 0.14<sup>n.s.</sup> | 2.91<sup>·</sup> | 0.00<sup>n.s.</sup> | 2.49<sup>·</sup> | 2.23<sup>n.s.</sup> | 0.46<sup>n.s.</sup> |
| CII | 142 | 2.37<sup>n.s.</sup> | 1.36<sup>n.s.</sup> | 0.07<sup>n.s.</sup> | 2.41<sup>n.s.</sup> | 0.02<sup>n.s.</sup> | 3.16<sup>*</sup> | 1.90<sup>n.s.</sup> | 0.45<sup>n.s.</sup> |
| State 4<sub>o</sub> | 144 | 9.82<sup>**</sup> | 4.59<sup>*</sup> | 0.65<sup>n.s.</sup> | 1.76<sup>n.s.</sup> | 3.28<sup>·</sup> | 2.57<sup>·</sup> | 0.70<sup>n.s.</sup> | 1.27<sup>n.s.</sup> |
| CIV capacities | 130 | 2.85<sup>·</sup> | 1.48<sup>n.s.</sup> | 1.00<sup>n.s.</sup> | 2.75<sup>·</sup> | 0.19<sup>n.s.</sup> | 1.80<sup>n.s.</sup> | 1.11<sup>n.s.</sup> | 1.14<sup>n.s.</sup> |

· p<0.1; \* p<0.05; \*\* p<0.01; \*\*\* p<0.001

### 5.1. Effects of acute *in vitro* warming on mitochondrial function in cold-conditioned fish

Acute warming of heart muscle fibers of cold-conditioned fish led to slightly but not significantly increased respiration rates. OXPHOS (Figure 1), OXPHOS-CI (Figure 2) and OXPHOS-CII (Figure 3) respiration rates tended to increase with acute warming, however, this was only significant for OXPHOS-CII in the Δ1000 group (LME, p <0.05). The relative contributions of CI and CII to OXPHOS respiration did not differ between conditions (CI-fraction: 53.4±3.3 – 61.1±3.3 %, Figure S1 and CII-fraction: 38.8±3.3 – 46.5±3.3 %, Figure S2).

**Figure 1:**
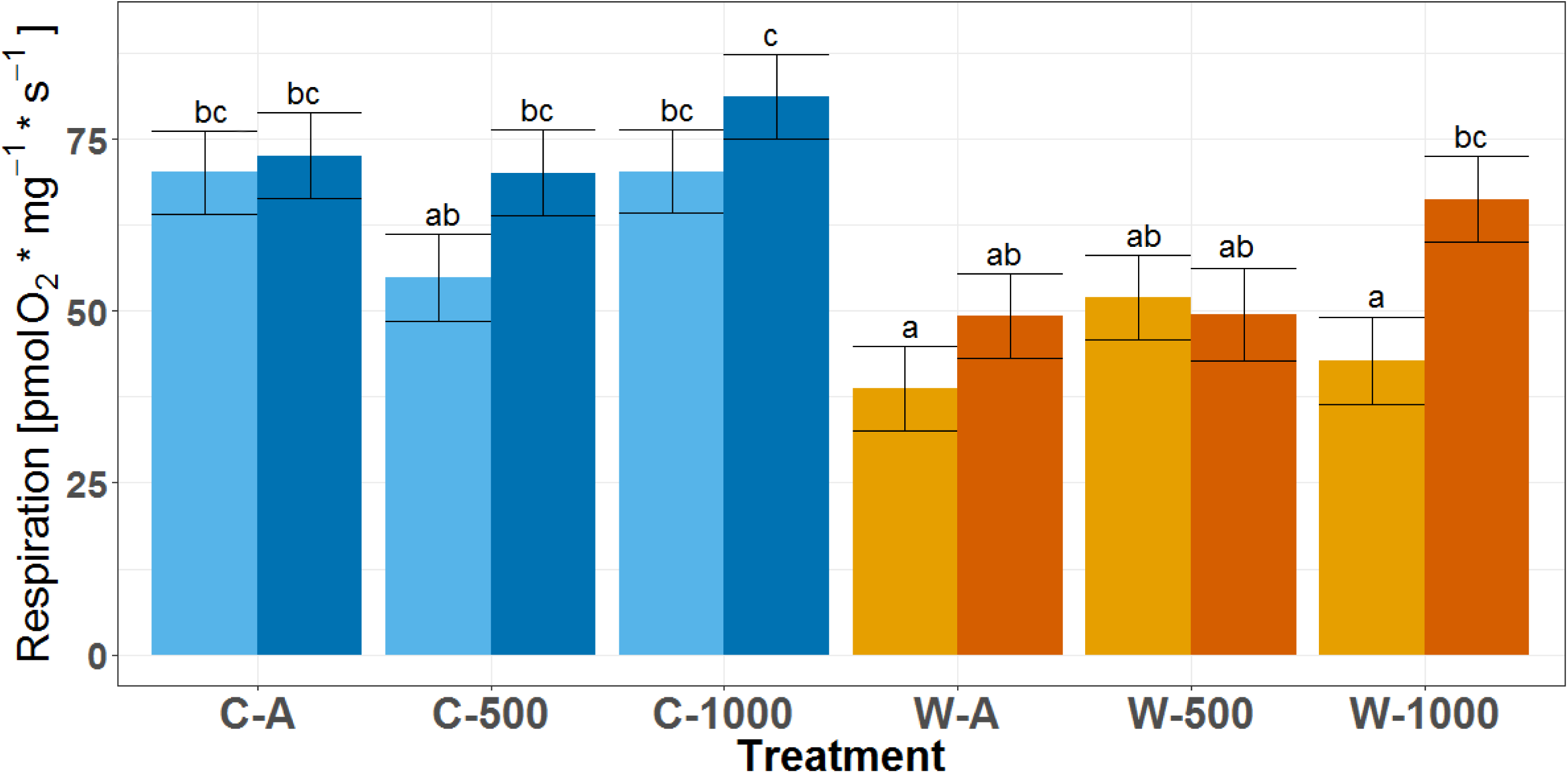
OXPHOS respiration rates of permeabilized heart fibers of European seabass. Shown are lsmeans ± s.e.m. Different letters indicate significant differences (LME, p<0.05); blue – cold life conditioned fish (C), orange – warm life conditioned fish (W), light color – cold assay temperature, dark color – warm assay temperature, A – Ambient *P*CO_2_, 500 – ambient + 500 µatm CO_2_, 1000 – ambient + 1000 µatm CO_2_; n=11-18.

**Figure 2:**
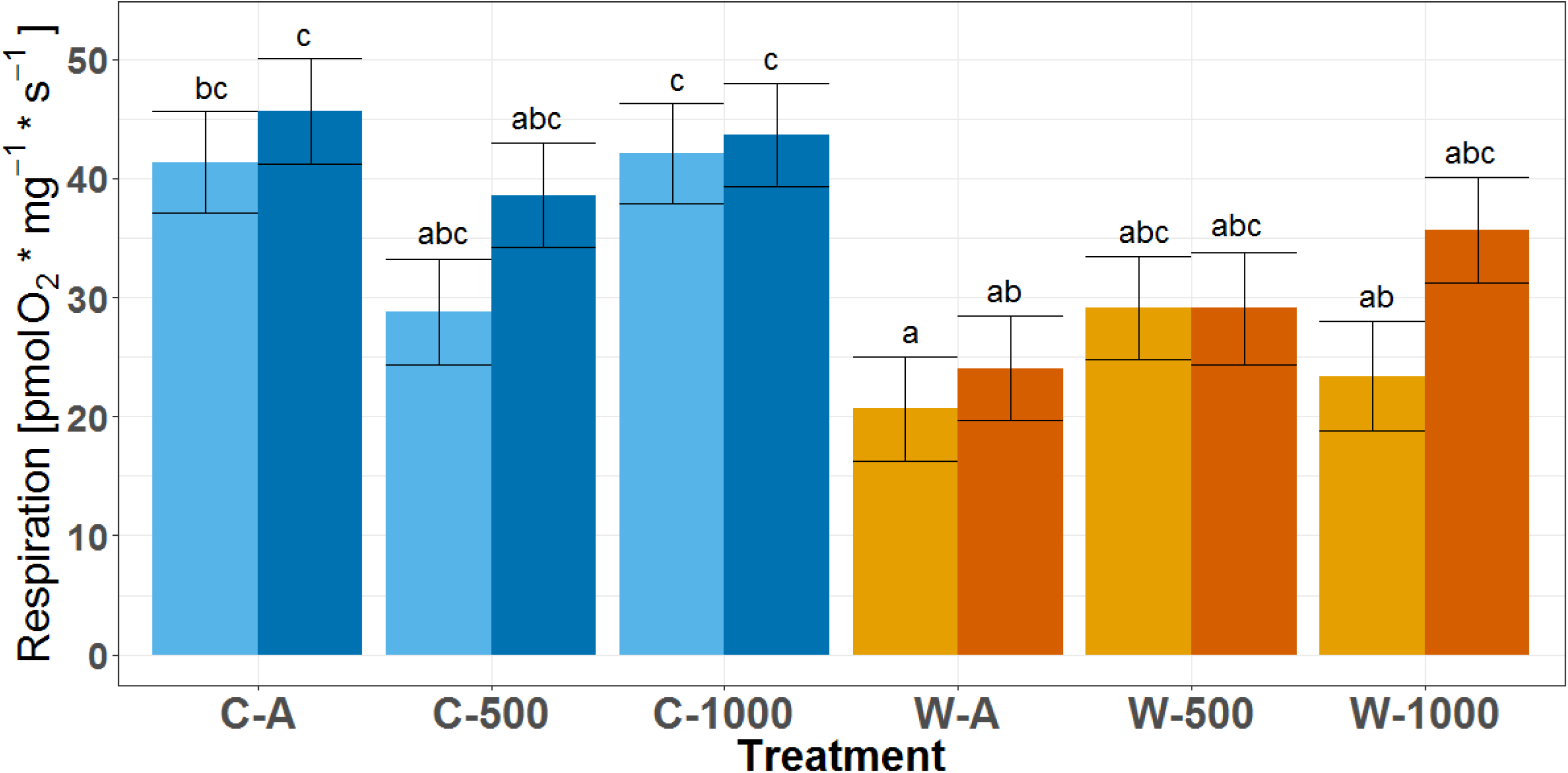
State 3 complex I (CI) respiration rates of permeabilized heart fibers of European seabass. Shown are lsmeans ± s.e.m. Different letters indicate significant differences (LME, p<0.05); blue – cold-conditioned fish (C), orange – warm-conditioned fish (W), light color – cold assay temperature, dark color – warm assay temperature, A – Ambient *P*CO_2_, 500 – ambient + 500 µatm CO_2_, 1000 – ambient + 1000 µatm CO_2_; n=11-18.

**Figure 3:**
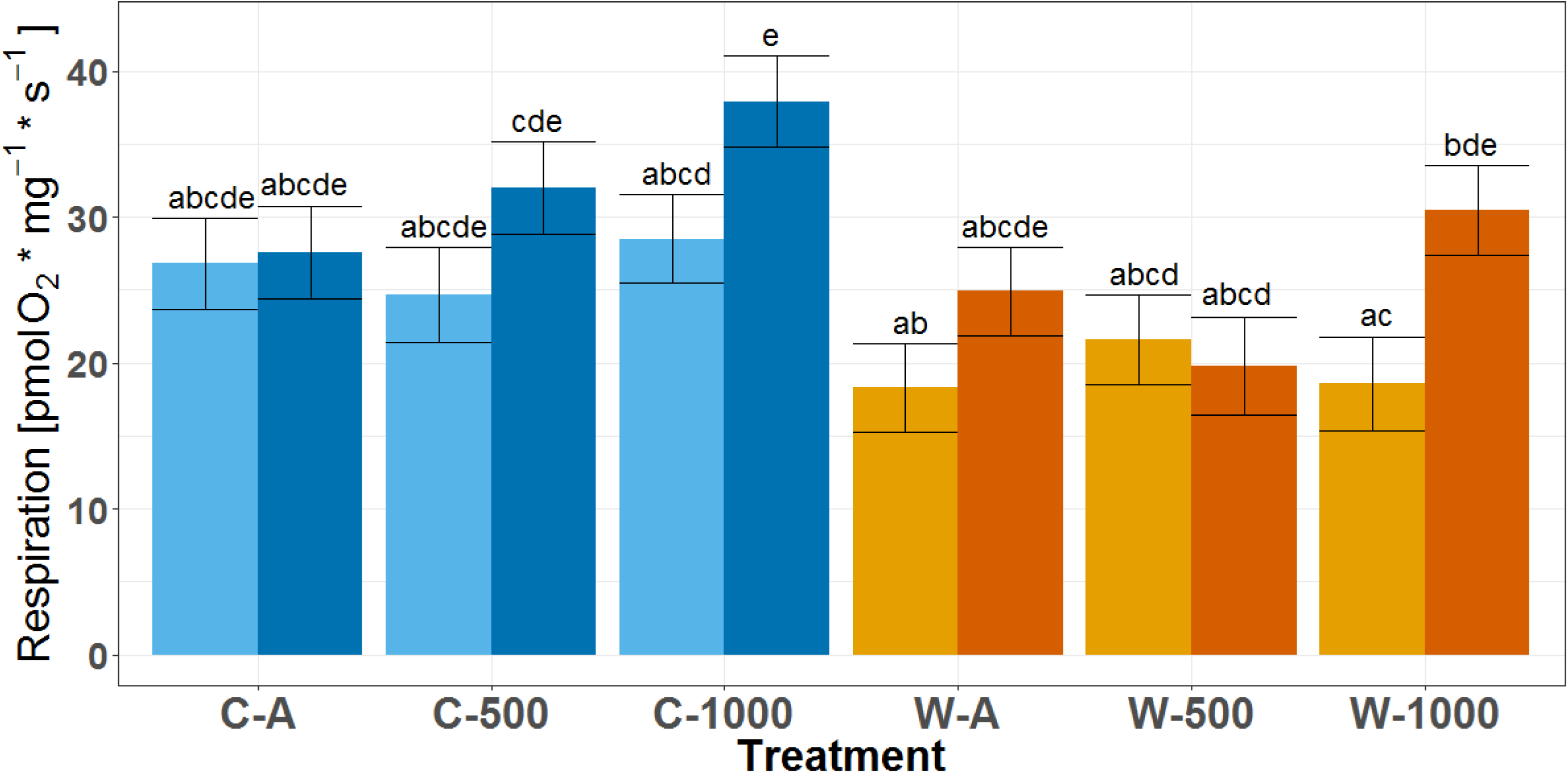
State 3 complex II (CII) respiration rates of permeabilized heart fibers of European seabass. Shown are lsmeans ± s.e.m. Different letters indicate significant differences (LME, p<0.05); blue – cold-conditioned fish (C), orange – warm-conditioned fish (W), light color – cold assay temperature, dark color – warm assay temperature, A – Ambient *P*CO_2_, 500 – ambient + 500 µatm CO_2_, 1000 – ambient + 1000 µatm CO_2_; n=11-18.

In CIV, the increase in respiration in response to acute warming was more pronounced, and significant in both hypercapnia groups (LME, Δ500: p<0.05 and Δ1000: p<0.05) but not in fish at ambient *P*CO_2_ (Figure 4). As OXPHOS and CIV respiration rates were affected similarly, CIV capacities did not change with acute warming (Figure S3).

**Figure 4:**
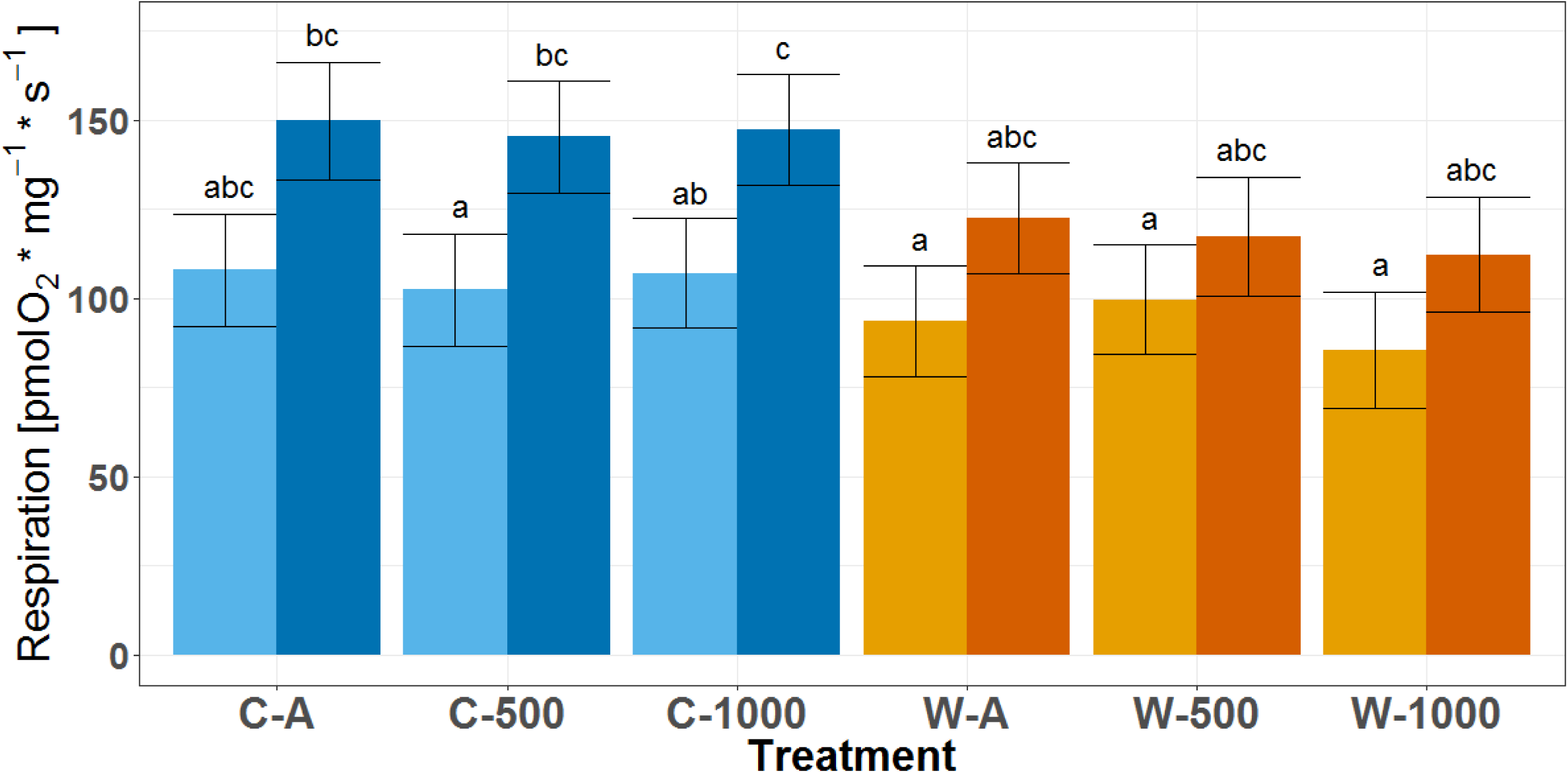
Complex IV (CIV) respiration rates of permeabilized heart fibers of European seabass. Shown are lsmeans ± s.e.m. Different letters indicate significant differences (LME, p<0.05); blue – cold-conditioned fish (C), orange – warm-conditioned fish (W), light color – cold assay temperature, dark color – warm assay temperature, A – Ambient *P*CO_2_, 500 – ambient + 500 µatm CO2, 1000 – ambient + 1000 µatm CO_2_; n=11-18.

The increase of state 4_o_ respiration rates with acute warming was significant in the A and Δ1000 treatments (LME, p<0.05 and p<0.01, respectively; Figure 5). Although OXPHOS and state 4_o_ respiration rates seemed to be affected similarly by acute warming, the increase in respiration rates was more pronounced in state 4_o_ than in OXPHOS, resulting in significant effects of assay and acclimation temperature on state 4_o_-fraction (Table 3). Despite these significances in the LME model, no significant differences were visible in the posthoc results (Figure S4). However, as no effect of *P*CO_2_ alone or in interaction with acclimation or assay temperature was detected by the model, we pooled data over *P*CO_2_ treatments (Table 4), which emphasized that acute warming lead to impaired mitochondria in the cold-conditioned fish, indicated by a significantly higher state 4_o_-fraction in warm compared to cold assay temperature (LME, p<0.05).

**Table 4:** Fraction of state 4_o_ respiration on OXPHOS respiration (State 4_o_-fraction) of permeabilized heart fibers of European seabass. Shown are lsmeans ± s.e.m. pooled over all *P*CO_2_ conditions, different letters indicate significant differences (LME, p<0.05). CL – cold-conditioned fish, WL – warm-conditioned fish, AT – assay temperature, n=43-48.

| AT | CL [%] |  |  | WL [%] |  |  |
| --- | --- | --- | --- | --- | --- | --- |
| Cold | 16.58 | ± | 1.60<sup>a</sup> | 14.58 | ± | 1.65<sup>a</sup> |
| Warm | 20.79 | ± | 1.65<sup>b</sup> | 14.41 | ± | 1.70<sup>a</sup> |

**Figure 5:**
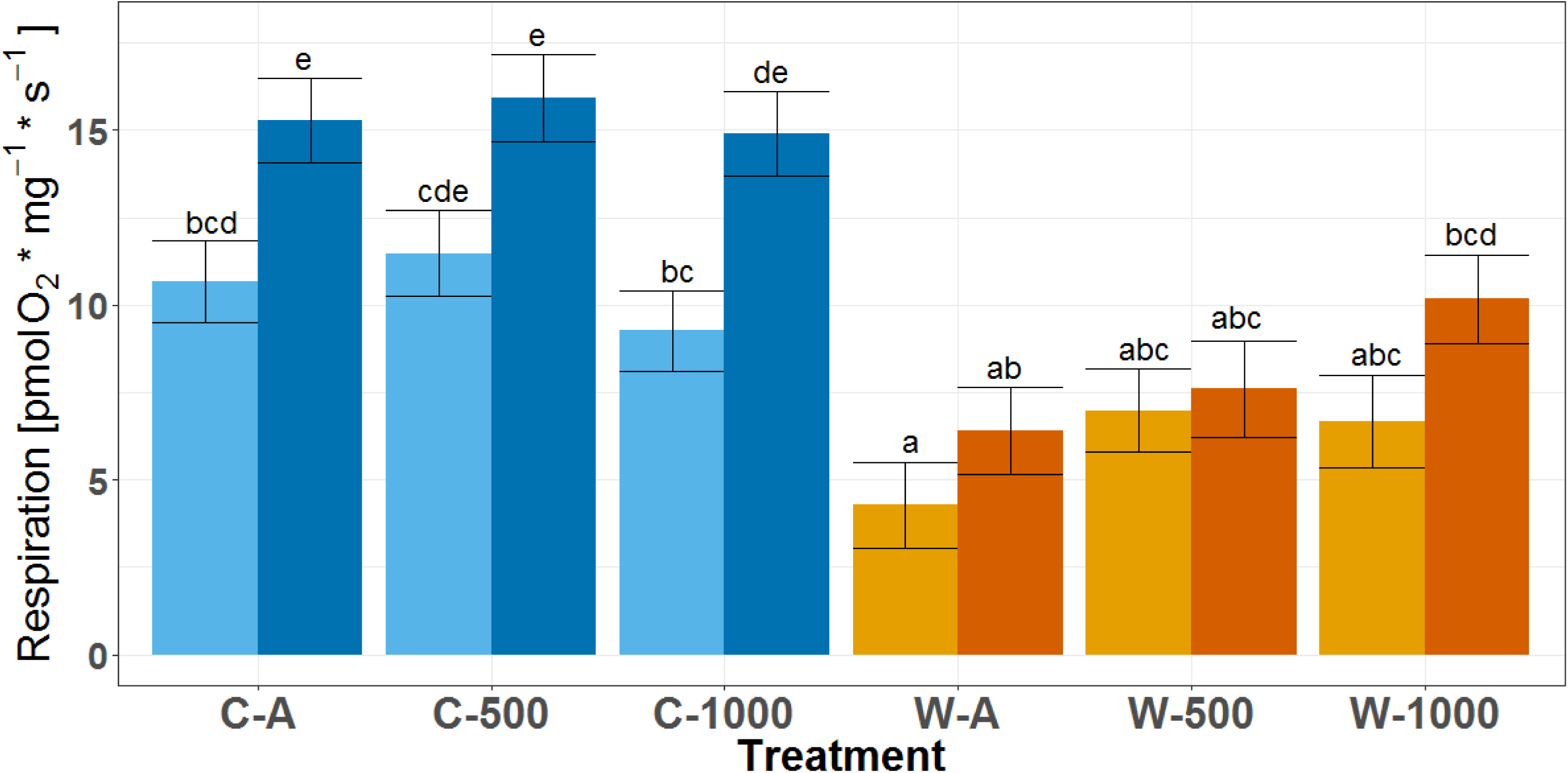
State 4o respiration rates of permeabilized heart fibers of European seabass. Shown are lsmeans ± s.e.m. Different letters indicate significant differences (LME, p<0.05); blue – cold life condition (C), orange – warm life condition (W), light color – cold assay temperature, dark color – warm assay temperature, A – Ambient *P*CO_2_, 500 – ambient + 500 µatm CO_2_, 1000 – ambient + 1000 µatm CO_2_; n=11-18.

### 5.2. Effects of acute *in vitro* cooling on mitochondrial function in warm-conditioned fish

In mitochondria of the warm-conditioned fish no general pattern with acute cooling *in vitro* could be observed. Significant differences were only found in the Δ1000 treatment, were acute cooling led to significantly decreased OXPHOS and OXPHOS-CII respiration rates (LME, p<0.01, Figure 1 and p<0.01, Figure 3, respectively). OXPHOS-CI (Figure 2), CIV (Figure 4) and state 4_o_ (Figure 5) respiration rates were not affected by acute cooling. CI fraction (50.2±3.2 – 57.5±3.3 %, Figure S1), CII-fraction (41.7±3.2 – 49.7±3.1 %, Figure S2), state 4_o_ fraction (Figure S4) and CIV capacities (Figure S3) were also not affected by acute cooling and remained stable over all conditions.

### 5.3. Effects of long-term acclimation to warmer temperatures on mitochondrial function

Acclimation to warmer temperatures lead to decreased respiration rates in permeabilized heart fibers. This was significant for OXPHOS respiration rates in the A and Δ1000 treatments at cold assay temperature (Figure 1, LME, p<0.001 and p<0.01, respectively), as well as in OXPHOS-CI in A (Figure 2, LME, both assay temperatures: p<0.01) and Δ1000 (LME, cold assay temperature, p<0.05). Consequently, as both were affected similarly, no significant effect of acclimation temperature on CI-fraction was found.

In contrast, OXPHOS-CII was not significantly different in permeabilized heart fibers of warm and cold-conditioned fish. Although the LME model showed a significant interaction between acclimation temperature and *P*CO_2_ for CII-fraction, see Table 3, the posthoc test did not reveal any significant differences between individual treatments. Even when pooled over assay temperatures, no significant differences were found (data not shown).

CIV respiration rates as well as CIV capacities did not differ between warm-and cold-conditioned juveniles. In general, CIV capacities were 1.5 to 2 times higher than OXPHOS capacities, see Figure S3.

State 4_o_ respiration rates were decreased in warm-conditioned fish compared to cold-conditioned fish, although only significant in A (Figure 5, LME, p<0.01 and p<0.0001 for cold and warm assay temperatures, respectively) and in Δ500 fish (LME, p<0.0001 for warm assay temperature). When compared at their respective incubation temperatures, no significant differences between the temperature acclimation conditions could be observed.

Although the pattern of changes with life conditioning to warmer temperatures was similar in OXPHOS and state 4_o_ respiration rates, these changes were much more pronounced in state 4_o_ respiration rates, resulting in significantly decreased state 4_o_-fraction in warm-conditioned fish, see Table 3 and Table 4. Even though acclimation temperature was significant in the LME model no significant differences were visible in the posthoc results (Figure S4). The data pooled over *P*CO_2_ treatments show that at warm assay temperature, the cold-conditioned fish have a significantly higher state 4_o_-fraction then the warm-conditioned fish at warm assay temperature (LME, p<0.05). However, no significant difference was observed between the two acclimation temperatures if compared at the respective acclimation temperature or at cold assay temperature (Table 4).

During our experiments, mitochondria were well coupled in all treatments (RCR_o_ > 4), with significantly higher RCR_o_ in warm-conditioned fish than in cold-conditioned fish (LME, 9.43±1.38 and 6.46±1.33 respectively, p<0.05). There were no significant effects of assay temperature, *P*CO_2_ or any interaction terms on RCR_o_ (Table 3).

### 5.4. Effects of acclimation to different *P*CO_2_ on mitochondrial function

Elevated *P*CO_2_ alone did not have significant effect on any of the studied complexes and processes of the electron transport chain (Table 3). However, we found synergistic effects with temperature which became visible as interaction effects with acclimation or assay temperature only in the Δ500 or Δ1000 fish, as specified above.

## 6. Discussion

Mitochondrial functional capacities were examined in seabass juveniles raised in six combinations of three *P*CO_2_ and two temperature treatments. The data provide evidence that heart mitochondria of juvenile seabass are highly impaired by acute warming, as e.g. observed in increased state 4_o_ respiration rates. On the other hand, acclimation to warmer temperatures increased mitochondrial capacities in warm-conditioned fish in comparison to cold-conditioned fish, as seen through increased RCR_o_. OA did not severely affect mitochondrial functioning in juvenile seabass, indicated by no significant effects of *P*CO_2_ alone on mitochondrial capacities. However, OA intensified the effects of acute or long-term warming. This was most prominent in the high acidification warm life condition treatment, e.g. OXPHOS respiration rates were only significantly affected by acute temperature change in the W-Δ1000 fish. We observed the same effect in OXPHOS-CII (decrease due to acute cooling in W-Δ1000 fish), therefore the decrease in OXPHOS respiration rates during acute cooling were probably due to the decrease of OXPHOS-CII. CI, CIV and RCR_o_ were not affected by *P*CO_2_. This observation reflects the findings of previous studies in polar fish, where thermal effects on mitochondrial capacities were much more prominent than those of OA (e.g. Leo, et al., 2017; Strobel, et al., 2013). However, the reduced capacities of CII to cope with acute temperature changes in the W-Δ1000 fish is in agreement with other studies which found that CII was inhibited by elevated *P*CO_2_ in mammals and fish (Simpson, 1967; Wanders, et al., 1983; Strobel, et al., 2013). In juvenile seabass, CII was only inhibited by high *P*CO_2_ in warm-conditioned fish faced with an acute temperature decrease. Therefore juvenile seabass mitochondria appear in general able to cope with the inhibiting effect of high *P*CO_2_ on CII. Other studies suggested that mitochondria could employ anaplerotic mechanisms, such as decarboxylation of dicarboxylic acids (aspartic and glutamic acid), to stimulate CII with additional substrates to overcome inhibitory effects of high *P*CO_2_ (Langenbuch & Pörtner, 2002; Strobel, et al., 2013). This might have happened in mitochondria of juvenile seabass. However, if temperature or *P*CO_2_ conditions worsen, the decreased OXPHOS respiration rates due to reduced CII respiration could already occur at the respective acclimation temperature and not only under acute temperature change.

Acute warming impairs heart mitochondria of cold-conditioned juvenile seabass, by increasing state 4_o_ respiration rates. Although the slightly increased OXPHOS respiration rates after acute temperature increase would indicate a potential for higher mitochondrial capacities, state 4_o_ respiration rates increased significantly in the C-A and C-Δ1000 groups under acute warming. As state 4_o_ respiration rates increased more than OXPHOS respiration rates, the state 4_o_-fraction increased as a consequence, pointing at impaired mitochondrial capacities under acute warming. Another indicator for mitochondrial functionality is the respiratory control ratio (RCR_o_), which is calculated from OXPHOS and state 4_o_ respiration rates. Although OXPHOS respiration rates increased not as much with acute warming than state 4_o_ respiration rates, RCR_o_ values were not yet decreased. However, increased state 4_o_ respiration rates as well as increased state 4_o_-fraction indicate decreasing mitochondrial membrane integrity, which translates to mitochondria producing less ATP for the same amount of oxygen consumed. Increases in mitochondrial enzyme activity and respiration rates of mitochondrial complexes, as well as OXPHOS respiration rates with acute temperature increase had been shown in other fish species before, e.g. Antarctic nototheniids, European perch and Atlantic cod (Strobel, et al., 2013; Ekström, et al., 2017; Leo, et al., 2017). However, Iftikar and Hickey (2013) showed in hearts of New Zealand triple fin fishes that compromised mitochondria at acutely elevated temperature will ultimately lead to heart failure. Consequently, as acute warming of only 5°C impaired mitochondria of the cold-conditioned fish, we could conclude from our findings that cold-conditioned juvenile seabass might not be able to cope with large acute temperature changes due to heart failure. This reduced tolerance to acute temperature increase in the cold-conditioned fish seems to contradict the fact that European seabass are generally highly tolerant to a wide range of temperature (Via, et al., 1998; Claireaux & Lagardère, 1999). It also seems to contradict the high critical thermal maximum (CT_max_) of European seabass (28.12±0.09°C to 32.50±0.04°C in Mauduit, et al., 2016; and 31.3±0.3°C in Anttila, et al., 2017). However, in the study of Antilla et al. (2017) although CT_max_ was as higher than 30°C (fish acclimated at 17°C), arrhythmias already occurred at around 22°C. Furthermore, Arrhenius breakpoint temperature for maximum heart rate, which is connected to the thermal optimum of growth and aerobic scope, was 19.3±0.3°C (Anttila, et al., 2017). Additionally, the temperature with highest maximal heart rates, a measure for thermal limits of cardiac function, was 21.8±0.4°C in fish acclimated at 17°C (Anttila, et al., 2017). These findings fit well with the impaired mitochondria of acutely warmed cold acclimated fish and indicate that sublethal physiological limitations appear much below the critical thermal maxima. Furthermore, they support our conclusion that cold-conditioned juvenile sea bass might not be able to cope with large acute temperature changes, as they would suffer from heart failure at relatively low temperatures.

While acute warming impairs the performance of juvenile seabass heart mitochondria, comparison of cold and warm-conditioned fish indicates that the chosen cold acclimation temperature for this study is not the optimal temperature for Atlantic juvenile seabass, as the warm-conditioned fish showed higher mitochondrial capacities. The thermal biology of *D. labrax* had already been topic of several studies, although mainly on Mediterranean populations (e.g. Marangos, et al., 1986; Koumoundouros, et al., 2001; Lanari, et al., 2002; Person-Le Ruyet, et al., 2004; Dülger, et al., 2012), but see Russel et al. (1996), Ayala et al. (2003) and Gourtay et al. (2018). These studies found that hatching can occur from 8 to 20°C (Marangos, et al., 1986) and they determined optimal temperatures of 15 to 17°C for larval development (Koumoundouros, et al., 2001; Ayala, et al., 2003), 22 to 28°C for juvenile growth (Lanari, et al., 2002; Person-Le Ruyet, et al., 2004; Dülger, et al., 2012) and 22°C for aerobic scope and active metabolic rates (Claireaux & Lagardère, 1999, experiments were done on hybrids of different populations (personal observation)). However, in contrast to the Mediterranean populations, which are exposed to higher habitat temperatures (typical range from 13-29°C between winter and summer, Person-Le Ruyet, et al., 2004); the Atlantic population, from the coasts of France up to the North sea, experiences temperatures lower than 15°C for most of the year and mainly within the range of 6-18°C (Russel, et al., 1996). While there are genetically distinct populations in the Mediterranean (Bahri-Sfar, et al., 2000), no differences between the Northern and Southern distribution in the Atlantic has been proven yet. However, the Western Mediterranean population is genetically distinct from the Atlantic population with a separation area at the Gibraltar Strait (Naciri, et al., 1999) the populations also showed different growth characteristics (Vandeputte, et al., 2014). Additionally, differences in response to rearing temperature during larval development has been found in a study comparing Mediterranean and Atlantic seabass (Ayala, et al., 2001). Our fish are offspring of fish caught in the Bay of Biscay. In these latitudes, spawning, egg development and larval hatching takes place at temperatures of 8 to 13°C (Jennings & Pawson, 1992) and later life stages experience mainly temperatures between 6 and 18°C (Russel, et al., 1996). Therefore, the temperature range we used for incubating the larvae in the cold life condition was slightly above the natural temperature regime of seabass larvae from the chosen distribution area. However for juvenile incubation the cold life condition temperature range of 15-18°C was well within the natural temperature regime during summer. The temperature of the warm-conditioned juveniles was consequently above the temperature range of their natural habitat in the Bay of Brest. Our study thus provides evidence that the seabass from the chosen population are not yet adapted to lower temperatures, as the warm-conditioned juveniles displayed much better mitochondrial capacities than the cold-conditioned animals.

State 4_o_ and OXPHOS respiration rates were both significantly decreased in warm-conditioned fish. Additionally, RCR_o_ were significantly increased. RCR_o_ values above 4 indicate well-coupled and non-damaged mitochondria (Brand & Nicholls, 2011). Although mitochondria were well-coupled in all treatments, the increased RCR_o_ indicated that mitochondria of warm-conditioned fish possess higher capacities to oxidize substrates and produce ATP (Brand & Nicholls, 2011). Increased RCR_o_ are probably due to differences in the strength of change of OXPHOS and state 4_o_ respiration, which results in more efficient mitochondria, while still reducing oxygen consumption rates. This increased mitochondrial efficiency, exemplified by higher RCR_o_, could translate into higher growth rates: Shama et al. (2014) found that lower OXPHOS capacities resulted in optimized metabolic rates that could generate higher scopes for growth in sticklebacks acclimated to warmer temperatures. Additionally, in brown trout lower food intake and failure to grow was correlated to high state 4_o_ respiration rates of liver and muscle mitochondria and lower mitochondrial coupling in muscle mitochondria (Salin, et al., 2016). In other words, individuals with more efficient oxidative phosphorylation tend to grow better than those with less efficient mitochondria. Our study reflects these findings, the warm-conditioned juveniles showed higher RCR_o_ and lower OXPHOS respiration rates, while being significantly bigger than the cold-conditioned fish, even when compared at equal age in degree days.

In our study, CIV or cytochrome c oxidase (CCO) activities were not affected by life condition temperature. In the literature, contrasting results of CCO activities in European seabass can be found: Nathanailides et al. (2010) observed increased CCO activities with decreasing seasonal acclimation temperature, while Trigari et al. (1992) found decreased CCO activities with decreasing seasonal acclimation. As terminal electron acceptor of the electron transport system, CCO is important in aerobic respiration and was found to be the controlling site of mitochondrial respiration and ATP synthesis (Villani & Attardi, 2001; Gnaiger, 2009; 2012; Kadenbach, et al., 2010). CCO generally displays excess capacities, especially in heart tissue (Gnaiger, et al., 1998). In our study, CIV had excess capacities 1.5-2 fold higher than OXPHOS respiration rates in all treatments; therefore CIV is not limiting the capacities of juvenile seabass mitochondria. Excess capacities of 1.5-2 are within the scope found for other fish species e.g. in New Zealand triplefin fish (1.5-3.2, Hilton, et al., 2010), New Zealand wrasse (1.8-2.7, Iftikar, et al., 2015), in brown trout (1.9-2.6, Salin, et al., 2016) and Northern killifish (1.5-2.2, Chung, et al., 2017).

## 7. Conclusion

Although we used specimens originating from a northern population of seabass for this study, the results altogether indicate that the mitochondrial metabolism still supports temperatures as found in Mediterranean specimens. Consequently, juvenile seabass in the North Atlantic might benefit from increased temperatures. They also possess (within the limits of this study) high capacities to cope with OA, although they are less pronounced under OAW. The results of this study indicate that juvenile European seabass will be able to survive in an acidifying and warming ocean, however, there are further bottlenecks that may constrict their survival in a future climate. Firstly other life stages, especially egg and larval stages might be more vulnerable to temperature changes and increased *P*CO_2_ and secondly, other important traits, such as behavior or reproductive capacities and phenology might be affected differently by OAW. Consequently other traits and life stages will be analyzed during further studies.

## 8. Acknowledgements

We thank Hanna Scheuffele for her support in measuring mitochondrial respiration and Karine Salin for her critical comments. We acknowledge the technicians and researchers from the Laboratory of Adaptation, and Nutrition of Fish for their support in animal welfare during the weekends.

## 11. Author contributions

SH, LC, NLB, GC and FCM contributed to the design and conception of the research. SH did the mitochondrial respiration measurements. SH and FCM conducted the data analysis and interpretation and wrote the manuscript. LC, NLB and GC contributed to rearing the fish under experimental conditions, as well as drafting the manuscript.

